# Influenza-induced oxidative stress sensitizes lung cells to bacterial toxin-mediated necroptosis

**DOI:** 10.1101/2020.02.20.957951

**Authors:** Norberto Gonzalez-Juarbe, Ashleigh N. Riegler, Alexander S. Jureka, Ryan P. Gilley, Jeffrey Brand, John E. Trombley, Ninecia R. Scott, Peter H. Dube, Chad M. Petit, Kevin S. Harrod, Carlos J. Orihuela

**Affiliations:** Department of Microbiology, The University of Alabama at Birmingham, Birmingham, Alabama 35294-2170, USA; Department of Biochemistry and Molecular Genetics, The University of Alabama at Birmingham, Birmingham, Alabama 35294-2170, USA; Department of Microbiology, Immunology and Molecular Genetics, The University of Texas Health Science Center at San Antonio, San Antonio, TX 78229; Department of Anesthesiology and Perioperative Medicine, The University of Alabama at Birmingham, Birmingham, Alabama 35294-2170, USA; Infectious Diseases and Genomic Medicine Group, J Craig Venter Institute, 9605 Medical Center Drive, Suite 150, Rockville, MD 20850

**Keywords:** Pneumonia, Influenza A virus, *Streptococcus pneumoniae*, epithelial cells, necroptosis, cell death, inflammation

## Abstract

**Rationale:** Pneumonia caused by Influenza A virus (IAV) co- and secondary bacterial infections are characterized by their severity. Previously we have shown that pore-forming toxin (PFT)-mediated necroptosis is a key driver of acute lung injury during bacterial pneumonia. Here, we evaluate the impact of IAV on PFT-induced acute lung injury during co- and secondary *Streptococcus pneumoniae* (*Spn*) infection.

**Objectives:** Determine the impact of IAV infection on bacterial PFT-mediated lung epithelial cell (LEC) necroptosis. Determine the molecular basis for increased sensitivity and if inhibition of necroptosis or oxidative stress blocks IAV sensitization of LEC to PFT.

**Methods:** Mice and cells were challenged with IAV followed by *Spn*. Necroptosis was monitored by measuring cell death at fixed time points post-infection and immunofluorescent detection of necroptosis. Wildtype mice and LEC were treated with necroptosis inhibitors. Necroptosis effector molecule MLKL deficiency was tested for infection synergy. Oxidative damage to DNA and lipids as result of infection was measured *in vitro* and *in vivo*. Necroptosis and anti-oxidant therapy efficacy to reduce disease severity was tested *in vivo*.

**Measurements and Main Results:** IAV synergistically sensitized LEC for PFT-mediated necroptosis *in vitro* and in murine models of *Spn* co-infection and secondary infection. Pharmacological induction of oxidative stress *sans* virus sensitized cells for PFT-mediated necroptosis. Necroptosis inhibition reduced disease severity during secondary bacterial infection.

**Conclusions:** IAV-induced oxidative stress sensitizes LEC for PFT-mediated necroptosis. This is a new molecular explanation for severe influenza-associated bacterial infections. Necroptosis inhibitors are potential therapeutic strategies to reduce IAV-primed bacterial pneumonia severity.

**Summary:** Here we demonstrate that Influenza A virus (IAV) infection synergistically sensitizes lung cells to bacterial pore-forming toxin (PFT)-mediated necroptosis. Moreover, this contributes to the severity of lung injury that is observed during co- and secondary infection with *Streptococcus pneumoniae*. IAV-induced oxidative stress was identified as a key factor contributing to cell sensitization and induction of oxidative stress *sans* virus was sufficient to synergistically enhance susceptibility to PFT-mediated killing. Our results advance our understanding on the molecular basis of co- and secondary bacterial infection to influenza and identifies necroptosis inhibition and antioxidant therapy as potential intervention strategies.

## INTRODUCTION

Influenza A virus (IAV) is the most common cause of human influenza (flu) (1), infecting 4-8% of the U.S. population annually (2). Worldwide, the World Health Organization estimates that flu affects approximately 1 billion individuals annually, with 3 to 5 million cases of severe disease and a resulting 300,000 to 500,000 deaths (3). While IAV alone is capable of considerable morbidity and mortality, clinical and molecular epidemiology have shown that the most serious infections are frequently associated with co-infections or a secondary infection with a bacterial pathogen. *Streptococcus pneumoniae* (*Spn*; the pneumococcus) is the leading cause of community-acquired pneumonia and by far the most common bacterium associated with IAV infections (4). Highlighting the seriousness of IAV/*Spn* co-infections, 34%-55% of the deaths linked to the 2009 IAV pandemic were associated with bacterial infections, with *Spn* the most common bacteria identified (5, 6).

Over the past 20 years a number of seminal discoveries have helped to explain, at the molecular level, the synergy observed during IAV/*Spn* super-infection. Key findings include the observation that IAV neuraminidase cleaves terminal sialic acid on host cell glycoconjugates exposing normally cryptic antigens for bacterial attachment (7). Viral neuraminidase-cleaved sialic acid serves as a nutrient for *Spn* and promotes bacterial outgrowth (8). IAV-induced down regulation of ion channels in bronchial epithelial cell results in dysregulated pulmonary fluid homeostasis that favors bacterial replication (9). Fever, cytokines, and alarmins released from IAV-infected dying cells elicit a transcriptional response from *Spn* that causes it to disperse from biofilms and enhances its virulence (10). In addition, IAV-induced interferon (IFN) gamma down regulates expression of scavenger receptors on macrophages, such as MARCO, that are required for uptake of *Spn* in absence of capsule specific antibody (11). Finally, the immune response induced by IAV is inappropriate for clearance of bacteria and enhances pulmonary injury (12, 13). It is noteworthy, that the majority of this work has not focused on events that occur within lung epithelial cells (LEC), which are the nexus of co-infection.

Necroptosis is a programmed form of cell death that results in host cell membrane failure, i.e. necrosis. It is inflammatory due to the release of cytoplasmic contents that serve as alarmins. Canonically, necroptosis is regulated by receptor-interacting serine-threonine kinase (RIPK)1, that activates RIPK3. Subsequently, RIPK1/RIPK3 activates the necroptosis effector molecule MLKL through phosphorylation, p-MLKL, which targets cell membranes leading to cell rupture and death (14, 15). Importantly, both IAV and bacterial pore-forming toxins (PFT), such as pneumolysin produced by *Spn*, have recently been shown to induce necroptosis of LEC (16–20). For IAV, this has been shown to be the result of viral RNA interactions with DAI (also known as Zbp or DLM-1), a sensor for cytoplasmic nucleic acid, which activates RIPK3. Necroptosis of virally infected LEC is thought to be beneficial as RIPK3 KO and MLKL/FADD double KO mice were considerably more susceptible to IAV (18). More recently, our research group has shown that membrane damage caused by the PFT of *Spn* and *Serratia marcescens* resulted in ion dysregulation which activated RIPK1 (21). In contrast to IAV infection, necroptosis during bacterial pneumonia was detrimental and exacerbated bacterial outgrowth, pulmonary injury, and loss of alveolar-capillary integrity (21, 22). Critically and up to this point, the role of necroptosis during IAV/bacteria co-infection was not known. Herein we determined its consequence and determined the underlying molecular mechanism responsible using *Spn* as the prototype bacterial pathogen.

## METHODS

### Ethics Statement

Animal experiments were approved by the Institutional Animal Care and Use Committee at The University of Alabama at Birmingham (Protocol # 20358). Human LEC were harvested from whole lung sections obtained from the International Institute for the Advancement of Medicine (23). The use of primary tissue, obtained in de-identified fashion, does not meet the criteria for human subject research.

### IAV and *Spn*

Pandemic H1N1 A/California/7/2009 (pdmH1N1) and H1N1 A/Puerto Rico/8/1934 (PR8) influenza viruses were propagated in MDCK cells. *Spn* serotype 4 strain TIGR4 and its derivatives were used for all studies (24). TIGR4 mutants deficient in *ply* (Δ*ply*), the gene encoding pneumolysin, and *spxB* (Δ*spxB*), the gene encoding pyruvate oxidase, have been described (25). We also used mutants provided by Dr. Jeffrey Weiser (New York University, NY). These were matched strains of TIGR4 (TIGR4_JW_), TIGR4 lacking pneumolysin, TIGR4_JW_ Δ*ply*), a TIGR4 point mutant deficient in pore formation (TIGR4_JW_ W433F), and a corrected mutant (TIGR4_JW_ *ply*+) (26); these were used as a set. Recombinant pneumolysin (rPly) was purified from *E. coli* (27). *Staphylococcus aureus* alpha-toxin was purchased (Sigma-Aldrich, St. Louis, MO).

### Animal strains and infections

Male and female 8-week-old C57BL/6 mice were obtained from Taconic Biosciences (Rensselaer, NY). MLKL KO mice were made available by Dr. Warren Alexander (Walter and Eliza Hall Institute of Medical Research Parkville, Victoria, Australia) (28). For IAV/*Spn* co-infection, 8-week-old C57BL/6 mice were intranasally challenged with 250 PFU PR8. Five days post-influenza challenge, mice received by forced aspiration 5×10^5^ CFU *Spn* (29). For studies involving *Spn* secondary infection, i.e. after viral clearance, mice were challenged with 250 PFU pdmH1N1 and ten days post-influenza, challenged with 10^3^ CFU *Spn*.

### Cell Infections

A549 type II alveolar epithelial cells (23), MH-S mouse alveolar macrophages (30), and primary normal human bronchiolar epithelial cells (23), were infected with IAV at MOI 2 for 2 hours, and subsequently challenged with *Spn* at an MOI 10 for 4 hours. The majority of chemical inhibitors were obtained from Sigma-Aldrich. Exceptions include necrosulfonamide (Tocris Bioscience, QL, UK), GSK’872 and Nec1s (BioVision, Milpitas, CA), oseltamivir carboxylate (MCE, Monmouth, NJ), TNFR inhibitor R-7050 and TNF-α inhibitor SPD-304 (Cayman Chemicals, Ann Arbor, MI) and Pimodivir (AdooQ Bioscience, Irvine, CA). Cells receiving inhibitors were treated continuously beginning 1-hour prior to IAV infection. Pimodivir treated cells received the drug 2-hours prior to IAV challenge. A549 cells deficient in MLKL have been previously described (16). Cell death was evaluated by detection of lactate dehydrogenase (LDH) in culture supernatants (22). The presence of reactive oxygen species (ROS) was measured with the H2-DCF assay (Thermo Fisher Scientific, Waltham, MA). Lipid peroxidation was detected with the lipid peroxidation malondialdehyde (MDA) assay (Abcam). Antibodies against 8-hydroxydeoxyguanosine, an oxidative stress-mediated DNA damage marker, and HNE-J, a lipid peroxidation marker, were purchased (Abcam).

### Histology and Microscopy

The methods used for tissue processing, sectioning, and immunofluorescent microscopy are described (17, 29, 31). Images were captured using a Zeiss AxioXam MRm Rev3 and/or MRc cameras attached to a Zeiss AxioImager Z1 epifluorescent microscope (Carl Zeiss, Thornwood, NY) or a Leica LMD6 with DFC3000G-1.3-megapixel monochrome camera (Leica Biosystems, Buffalo Grove, IL). TUNEL (Promega, Madison, WI) and Annexin V (Abcam, Cambridge, UK) staining was done per manufacturer’s instruction. Cleaved caspase-3 staining was done using anti-cleaved-caspase-3 antibody (Abcam). Mean fluorescent intensity and densitometry of immunoblots was measured using ImageJ (32).

### Immunoblots and ELISA

Western blots for MLKL (1:1000, #37705, Cell Signaling Technologies), p-MLKL (1:1000, #37333S, Cell Signaling Technologies) and cytoskeletal actin (1:10000, #A300-485A, Bethyl Laboratories Inc., Montgomery, TX), were done as previously described (33). ELISA-based measurements for IFN-β, IFN-α and TNF-α were done using kits from PBL Assay Science (Piscataway, NJ) and InvivoGen (San Diego, CA).

### Statistical analyses

For non-parametric multiple group analyses we used a Kruskal-Wallis H test with Dunn’s post-hoc analysis. For parametric grouped analyses we used ANOVA with Sidak’s post-hoc analysis. For data with a single independent factor of two groups we used a Mann-Whitney U test. Survival comparisons were assessed using Log-rank (Mantel-Cox) test. Asterisks denote the level of significance observed: * = P ≤ 0.05; ** = P ≤ 0.01; *** = P ≤ 0.001; **** = P ≤ 0.0001. Statistical analyses were calculated using Prism 8 (GraphPad Software: La Jolla, CA).

## RESULTS

### Necroptosis is synergistically increased during IAV/Spn co-infection

Using an established mouse model of co-infection (34, 35), we recapitulated the synergy known to occur between IAV and *Spn*. Briefly, we observed a >50-fold increase in the amount of *Spn* present in bronchoalveolar lavage fluid (BALF) and blood (**Fig. 1A, B**), as well as a significant decrease in time to death following IAV/*Spn* challenge versus *Spn* or IAV alone (**Fig. 1C**). Importantly, ongoing IAV infection synergistically enhanced the number of lung cells undergoing necroptosis after *Spn* challenge; necroptosis activity in frozen lung sections was inferred by immunofluorescent detection of phosphorylated MLKL (p-MLKL) (**Fig. 1D, E**).

**Figure 1:**
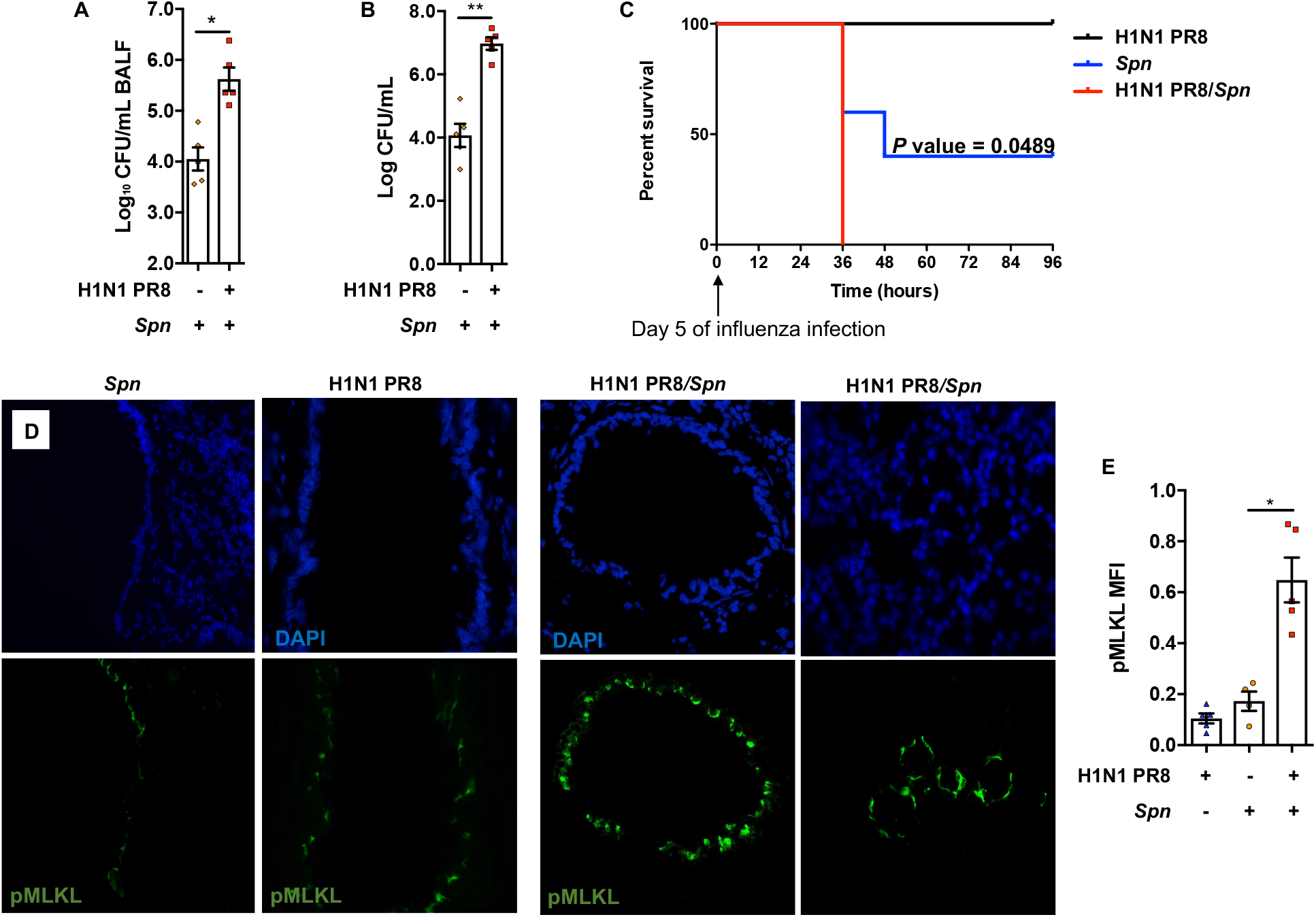
IAV/*Spn* co-infection leads to increased mortality and enhanced tissue necroptosis. 8-week-old C57Bl/6 mice were intranasally infected with H1N1 PR8 (250 PFU) for 5 days and subsequently challenged intratracheally with *Spn* strain TIGR4 at the LD_50_ dose of 5 x 10^5^ CFU. Mice were euthanized 24-hours post-secondary infection (n=4-5 mice/cohort). Bacterial titers in **A)** bronchoalveolar lavage (BALF) and **B)** blood of mice at time of sacrifice. **C)** Survival of mice challenged with IAV, *Spn*, or co-infected with *Spn* after 5 days of IAV (n=5); **D)** corresponding and representative images of frozen lung sections from infected mice immunofluorescent stained for p-MLKL (green) (n=4-5/cohort). **E)** Shown is the quantitation of p-MLKL levels in captured images calculated by mean fluorescent intensity.

To validate this *in vivo* observation and begin to dissect the molecular mechanisms underlying IAV-enhanced bacteria-induced necroptosis, we used an established *in vitro* co-infection model (36). Briefly, A549 type II alveolar epithelial cells were infected with either pdmH1N1 or PR8 at a MOI of 2 for two hours and then challenged with *Spn* at a MOI of 10 for another four hours. Importantly, A549 cytotoxicity was synergistic increased in cells challenged with both pathogens (**Fig. 2A, Fig. E1**). Similar results were also observed with MH-S murine alveolar macrophages (**Fig. E2**), indicating influenza-mediated sensitization to necroptosis is not restricted to airway epithelial cells. Of note, the enhanced death of A549 co-infected cells occurred without significant differences in bacterial titers versus control (**Fig. E3A**); indicating that the increased levels of necroptosis observed *in vivo* were not solely due to increased bacterial burden. Tumor necrosis factor (TNF) and IFN responses have been shown to promote necroptosis during viral infection (37). Along such lines, inhibition of TNF receptor 1 or blocking of TNF-α by pre-treatment of cells with R7050 or SPD304, respectively, did not reduce influenza-induced cell death potentiation in A549 cells *in vitro* (**Fig. E3B**). Moreover, the timeframe of the *in vitro* model did not lead to significant increases in the interferon response (**Fig. E3C**). Altogether, no evidence supporting a role for the synergistic initiation of receptor-mediated apoptosis was found *in vitro* or *in vivo* under the conditions tested (**Fig E4**).

**Figure 2:**
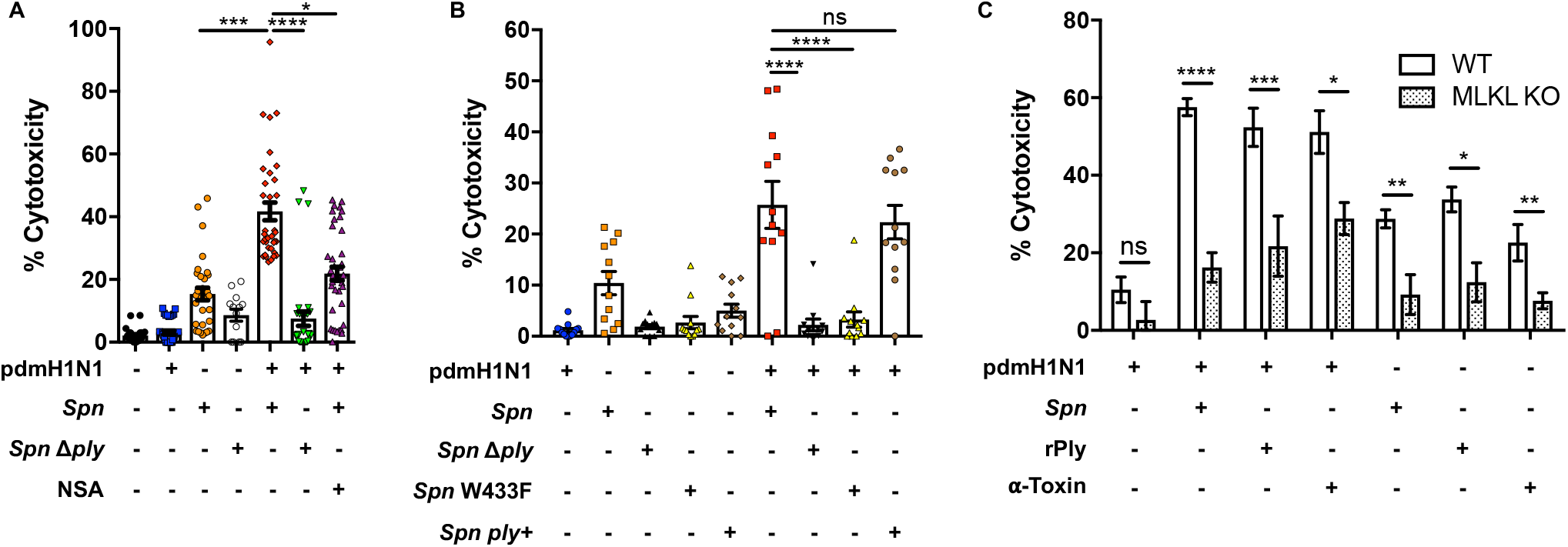
IAV infection promotes PFT-mediated cell death. **A)** LDH release was measured from A549 cells following infection with influenza A/California/7/2009 (pdmH1N1) at a MOI of 2 for 2 hours and challenge with wildtype *Spn* (*Spn*, in house strain) or Ply deficient derivative (*Spn* Δply) at an MOI of 10 for 4 additional hours. Cells were treated with necrosulfonamide (NSA, 10μM) when indicated. **B)** LDH cytotoxicity assay of supernatants from A549 cells was performed following infection with pdmH1N1 at an MOI of 2 for 2 hours and challenge with *Spn* strains and mutants obtained from Dr. Jeffrey Weiser at an MOI of 10 for 4 hours: *Spn* TIGR4 WT (*Spn*), Ply deficient mutant (*Spn* Δply), Ply point mutant deficient in pore formation (*Spn* W433F), and corrected mutant (*Spn ply*+). **C)** Cytotoxicity of A549 wildtype (white bars) or A549 MLKL deficient cells (dotted bars) was measured following the same challenge model as in panel a using *Spn*, recombinant pneumolysin (rPly), or alpha-toxin (α-Toxin).

### Pore-forming toxin activity is required for Spn-induced necroptosis during co-infection

*Spn*-mediated cytotoxicity of LEC was found to require the pore-forming activity of its PFT pneumolysin (**Fig 2A, B**). What is more, when A549 cells were treated with inhibitors of MLKL, necrosulfonamide (NSA) (**Fig 2A**) or RIPK3, GSK’ 872 (**Fig. E5**), the enhanced sensitivity of these cells to *Spn* killing was lost. Challenge of IAV-infected A549 cells with recombinant pneumolysin (rPly) or α-toxin (the PFT of *Staphylococcus aureus*, the second most common isolate during SBI to influenza (38)), recapitulated the potentiation of cell cytotoxicity observed with live bacterial infection (**Fig. E6**). Potentiation of necroptosis by IAV was confirmed by immunoblot and immunofluorescent staining which showed enhanced amounts of p-MLKL in A549 cells (**Fig. E7**). Further supporting a key role for necroptosis was the observation that A549 cells deficient in MLKL were protected against exacerbated PFT-mediated cell death after influenza infection (**Fig. 2C**). Moreover, that the same results were observed with primary normal human bronchiolar epithelial cells (nHBE) *ex vivo* (**Fig. E8**).

### IAV-induced oxidative stress sensitizes cells *in vitro* for PFT-mediated necroptosis

IAV-mediated oxidative stress has potent effects on pulmonary epithelial cells and the immune system (39). Therefore, it seemed plausible that the oxidative stress induced by the virus may be contributing towards the potentiation of pneumolysin-mediated necroptosis. In support of this notion, we observed that respiratory epithelial cells challenged *in vitro* with pdmH1N1or PR8 showed increased levels of lipid peroxidation (**Fig. 3A, Fig. E9A**) as measured by MDA and cellular ROS (**Fig. 3B, Fig. E9B**) as measured using H2-DCF. Importantly, and despite not having an effect on viral titers during the course of infection (**Fig. 3C**), pretreatment of A549 cells with the superoxide dismutase mimetic Tempol (40) prior to viral challenge reduced cell death and MLKL activation in co-infected cells (**Fig. 3D-E**). Directly implicating oxidative stress as a primer for PFT-induced necroptosis, treatment of cells with paraquat (22) enhanced the toxicity of rPly towards LEC and the observed potentiating effect of paraquat was abolished by treatment with Tempol (**Fig. 3F**). Identical results were observed using nHBE *ex vivo* (**Fig. 3G**) and replicated by addition of exogenous H_2_O_2_ in place of paraquat to A549 epithelial cells prior to rPly challenge (**Fig. E10**). Note that *Spn* also produces H_2_O_2_ via its metabolic enzyme SpxB (40). Yet, IAV potentiation of cell death was also observed in A549 cells challenged with *Spn* Δ*spxB* (**Fig. E11**), indicating that the priming effect of viral-induced ROS was sufficient. Importantly, inhibition of ROS in A549 cells with rotenone + thallium trifluoroacetate (mitochondria-dependent ROS inhibitor), apocynin (ADPH-dependent ROS inhibitor), allopurinol (xanthine oxidase-dependent ROS inhibitor) or mefenamic acid (cyclooxygenase-dependent ROS inhibitor) all conferred protection against death caused by co-infection (**Fig. 3H**). This results suggests that ROS potentiation of necroptosis may come from multiple cellular sources. Lastly, and to further probe the specificity of oxidative stress as a primer for necroptosis, we tested whether blockage of viral neuraminidase activity with oseltamivir (41) or treatment of cells with Pimodivir (VX-787) (42), a non-nucleoside polymerase basic protein 2 subunit inhibitor, impacted cell death. Neither of which did (**Fig. E12A**), despite the fact that viral titers were decreased by Pimodivir treatment (**Fig. E12B**).

**Figure 3:**
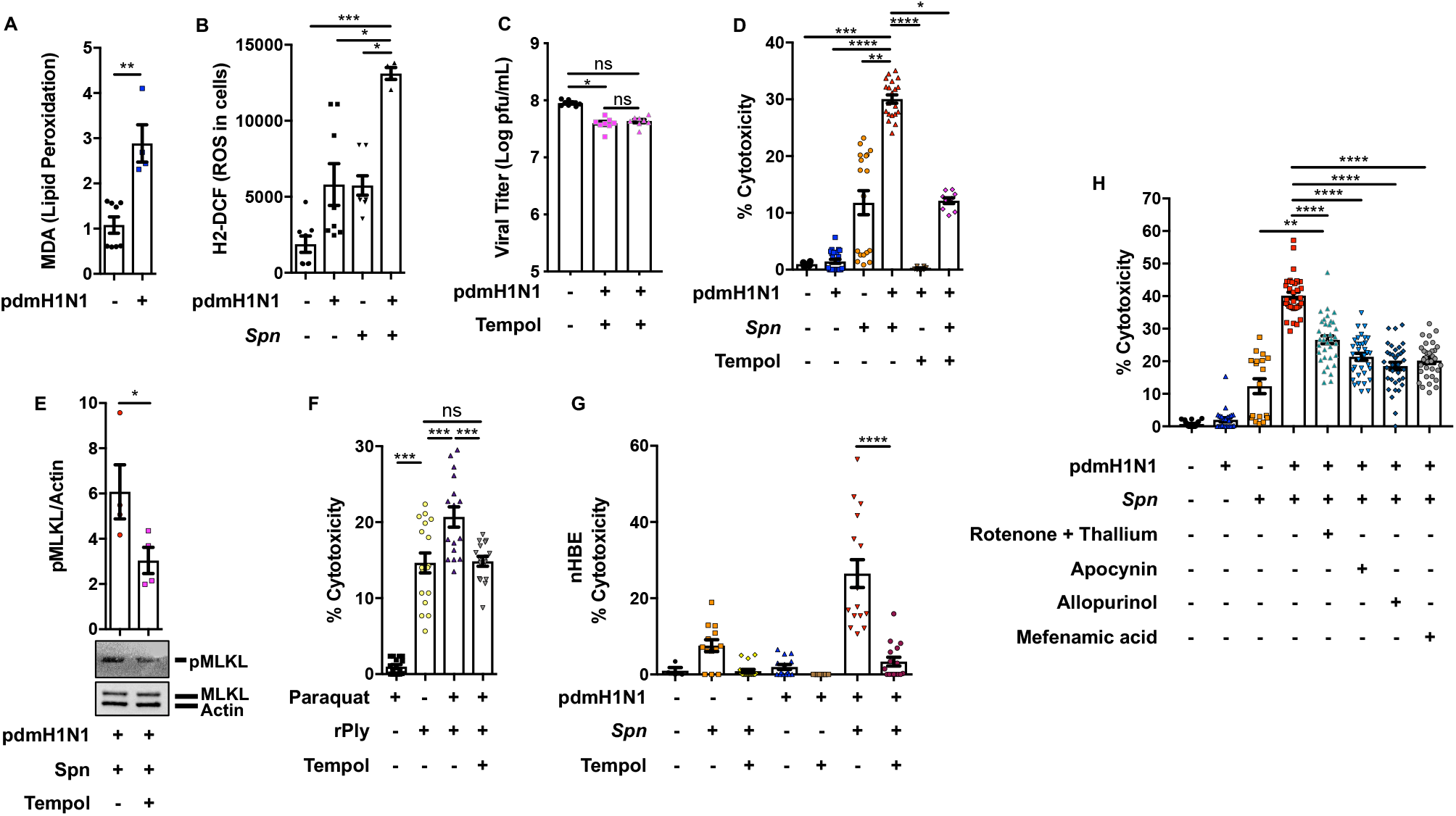
IAV-mediated oxidative stress potentiates pneumolysin-mediated necroptosis. **A)** Lipid peroxidation levels 4-hours after challenge with pdmH1N1 was measured by MDA. **B)** Levels of cellular ROS measured in A549 cells infected with pdmH1N1 at a MOI 2 for 2 hours then challenged with *Spn* at a MOI of 10 for 2 more hours. **C)** Viral titers quantified (Log PFU/mL) in A549 cells treated with Tempol (20μM) for 1-hour or 24-hours. **D)** Cytotoxicity and **E)** corresponding p-MLKL levels in A549 cells that were pre-treated with Tempol for 1-hour, infected with pdmH1N1 at a MOI 2 for 2 hours, then challenged with *Spn* at an MOI of 10 for 4 additional hours. **F)** Cytotoxicity was measured in A549 cells pre-treated with Tempol, then treated with Paraquat (10μM) for additional 2 hours, followed by challenge with rPly (0.1μg) for 2 hours. **G)** Cytotoxicity of *ex vivo* cultured primary normal human bronchial epithelial cells pre-treated with Tempol for 1-hour, infected with pdmH1N1 at a MOI 2 for 2 hours, and challenged with *Spn* at an MOI of 10 for 4 additional hours. **H)** LDH release from A549 cells pretreated with Rotenone + Thallium trifluoroacetate (10 nM/mL/10 nM/mL), a mitochondria-dependent ROS inhibitor; Apocynin (1μM/mL), a NADPH-dependent ROS inhibitor; Allopurinol (10nM/mL), a xanthine oxidase-dependent ROS inhibitor; and Mefenamic acid (20nM/mL), a cyclooxygenase-dependent ROS inhibitor following IAV and *Spn*, individually and together.

### IAV induced oxidative stress remains beyond viral clearance and maintains susceptibility to bacterial toxin mediated necroptosis

We examined whether residual oxidative stress induced by IAV helped to explain the enhanced susceptibility to bacterial infection that occurs even after IAV is cleared; i.e. in a secondary infection model. Lung sections from pdmH1N1-challenged mice 10 days post-IAV infection showed considerable evidence of oxidative damage to DNA of as well as lipid peroxidation (IF 8-Hydroxydeoxyguanosine and 4-Hydroxynonenal staining, respectively (43, 44)) in pulmonary tissue (**Fig. 4A-D**). Notably, these mice were confirmed to not have detectable virus (**Fig. 4E**). Similar to co-infection results (see Fig 1), if these mice were challenged with *Spn*, we observed a >100-fold increase in bacterial lung titers 2 days after *Spn* challenge (**Fig. 5A**). This was concomitant with greater lung consolidation, immune cell infiltration (**Fig. E13**), and substantially enhanced levels of lung necroptosis in co-infected mice versus those with *Spn* alone (**Fig. 5B-D**). Importantly, mice challenged with TIGR4 Δ*ply* in our secondary infection model had MLKL activation levels and bacterial titers equivalent to our negative control, i.e. mice infected with wildtype TIGR4 but also receiving the necroptosis inhibitor Nec-1s (**Fig. 5E-G, Fig. E14**). What is more, TIGR4 Δ*ply* challenged IAV-infected mice had decreased mortality versus controls (**Fig. 5H**). Interestingly, Tempol treatment at 12 and 24 hours post-*Spn* infection reduced the amount of necroptosis occurring in the airway in our secondary IAV/*Spn* infection model. This was despite not having an observed effect on *in vivo* levels of lipid oxidation (**Fig. 6A-E**). Tempol treatment also reduced bacterial burden within the airway of infected mice (**Fig. 6F**). Thus, necroptosis sensitizing ROS is primarily due to the virus, persisted beyond detectable IAV infection, and acted directly to sensitize the cell for necroptosis.

**Figure 4:**
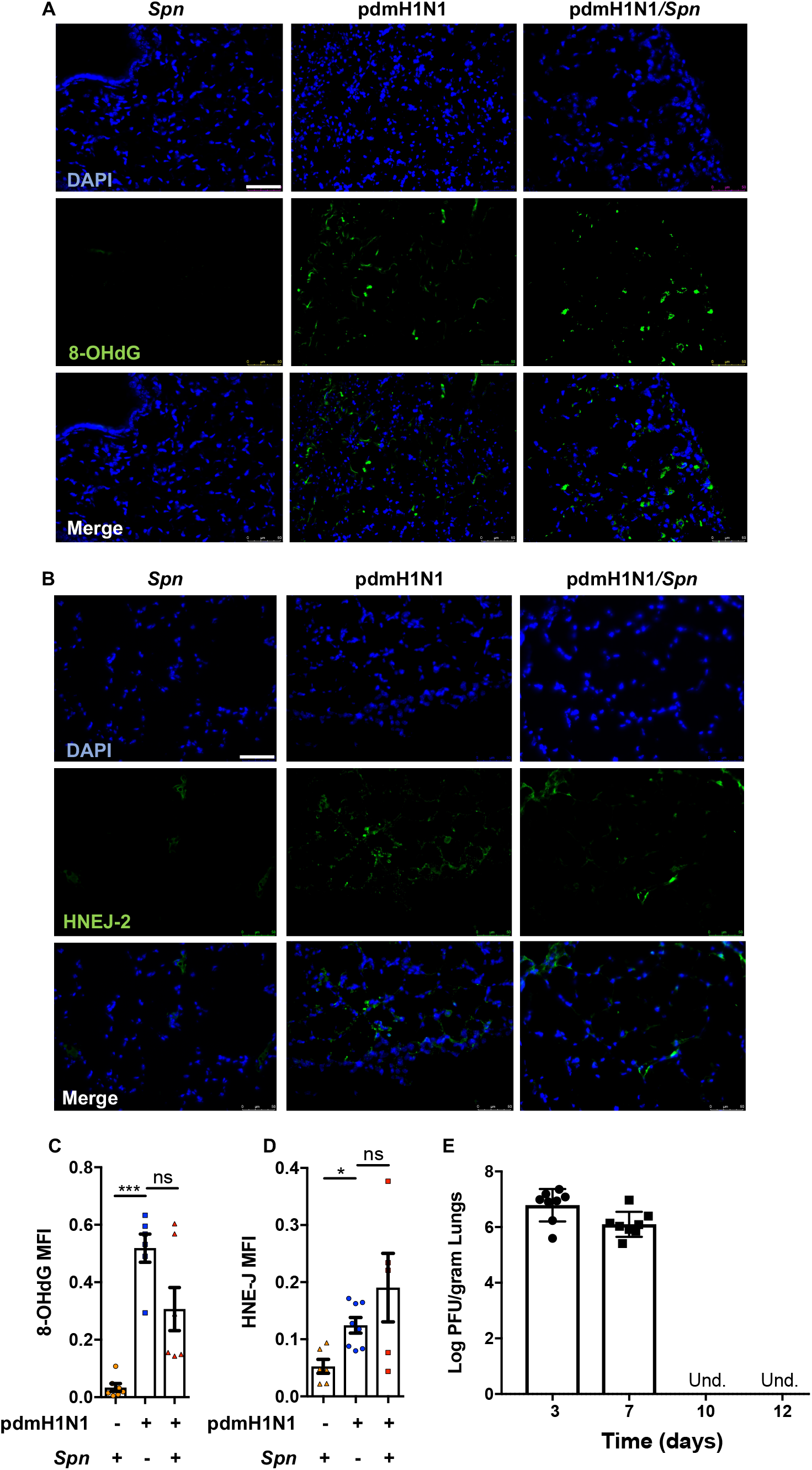
IAV-mediated oxidative stress persists after virus clearance *in vivo*. 8-week-old C57Bl/6 mice were intranasally infected with A/California/7/2009 (pdmH1N1) and 10 days later challenged intratracheally with *S. pneumoniae* (*Spn*). Mice were euthanized 48 hours after secondary infection (n=6-8 mice). Shown are representative immunofluorescent lung sections stained for **A)** 8-Hydroxydeoxyguanosine (8-OHdG) and **B)** 4-Hydroxynonenal (HNE-J). White bar denotes 50μm. **C-D)** Quantification of the mean fluorescent intensity (MFI) of 8-OHdG and HNE-J staining’s, respectively, was performed. **E)** Viral titers (Log PFU/gram) in lungs at days 3, 7, 10 and 12 post-IAV infection (n=8 mice per group) are indicated.

**Figure 5:**
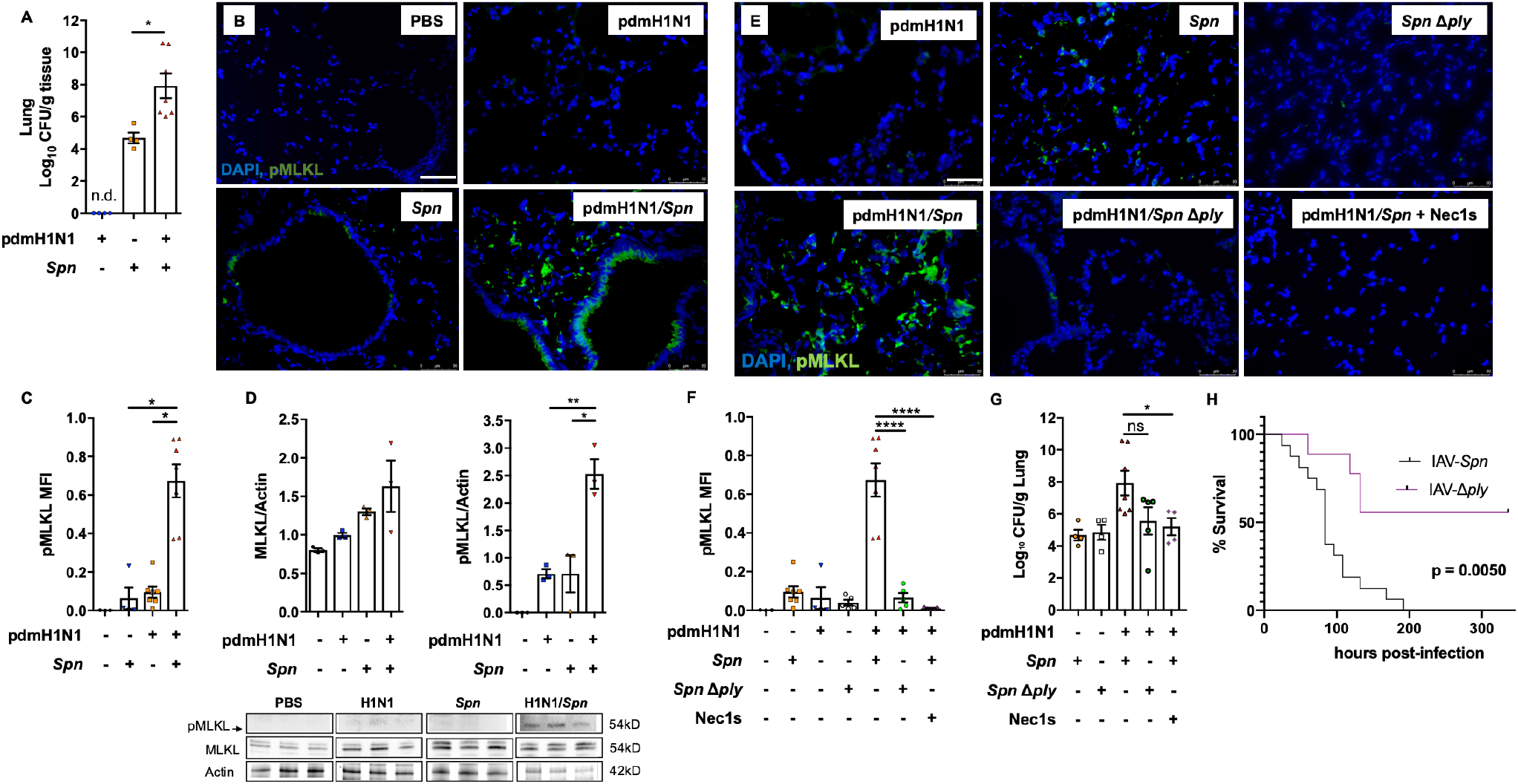
Influenza infection potentiates pneumolysin induced necroptosis activation during secondary *S. pneumoniae* challenge. 8-week-old C57Bl/6 mice were intranasally infected with A/California/7/2009 (pdmH1N1) and 10 days later challenged intratracheally with *S. pneumoniae* (*Spn*). Mice were euthanized 48 hours after secondary infection (n=3-7 mice). Shown are **A)** bacterial titers in homogenized lung samples, as well as **B)** representative images of corresponding lung sections stained for p-MLKL (3 sections stained per mouse). White bar denotes 50μm. **C)** Mean fluorescent intensity (MFI) for p-MLKL activity was measured. **D)** Densitometry and western blots for p-MLKL, MLKL and actin from mock, *Spn*, pdmH1N1 and pdmH1N1/*Spn* infected mice (n=3/cohort). **E-G)** 8-week-old C57Bl/6 mice were intranasally infected with A/California/7/2009 (pdmH1N1) and 10 days later challenged intratracheally with *Spn* or *Spn* Δply. Mice were euthanized 48 hours after secondary infection (n=4-7 mice). Treatment with Nec1s was done intraperitoneally at 12 and 24 hours following bacterial challenge. **E)** Shown are representative images of lung tissue sections stained for p-MLKL (separate points are average of 3 pictures per mouse) and **F)** mean fluorescent intensity of pMLKL staining. Corresponding **G)** lung bacterial titers (CFU/g tissue) calculated. **H)** Survival of C57Bl/6 mice following intranasal infection with pdmH1N1 for 10 days and subsequent intratracheal challenge with *Spn* or *Spn Δply* was monitored.

**Figure 6:**
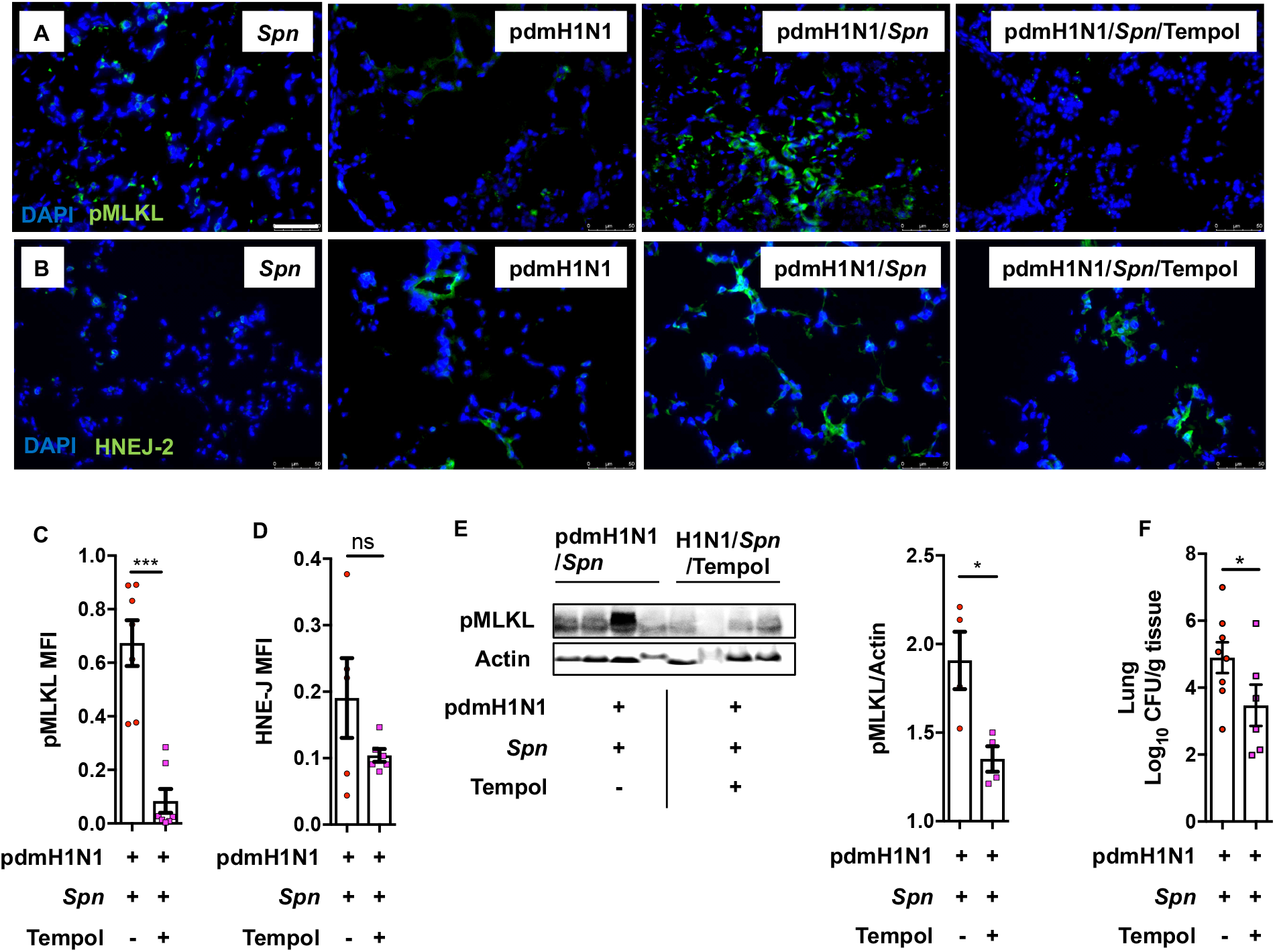
Therapeutic neutralization of ROS reduces necroptosis activation during secondary bacterial pneumonia. 8-week-old C57Bl/6 mice were intranasally infected with A/California/7/2009 (pdmH1N1) and 10 days later challenged intratracheally with *S. pneumoniae* (*Spn*). Mice were euthanized 48 hours after secondary infection (n=5-8 mice). Tempol treatment was done intraperitoneally at 12 and 24 hours post bacterial infection. Representative images of lung sections immunofluorescent stained for **A)** p-MLKL and **B)** 4 Hydroxynonenal (HNE-J). White bar denotes 50μm. **C-D)** Quantification of the mean fluorescent intensity (MFI) in corresponding captured images. **E)** Immunoblot for pMLKL and actin of pdmH1N1 infected mice, challenged with *Spn* with subsequent Tempol treatment and its densitometry quantification. **F)** Bacterial titers measured in lungs at time of death.

### *In vivo* necroptosis inhibition reduces severity of secondary bacterial infection to influenza

While no changes in oxidative stress induced DNA damage were observed (**Fig. E15**), MLKL deficient mice with secondary *Spn* infection had reduced bacterial titers, reduced lung consolidation, and a reduction in overall TUNEL positive staining in lung sections (a general marker of cell death) (**Fig. 7A-E**). In addition, lungs of MLKL KO Mice showed decreased levels of IFN-α and -β, suggesting a possible role for necroptosis in the IFN response during secondary infections (**Fig. 7F-G**). Most importantly, MLKL KO mice had greater survival versus control in our secondary infection model (**Fig. 7H**). Altogether, our results implicate oxidative-stress enhanced PFT-mediated necroptosis activity as a major driver of disease severity and lung injury during co- and secondary infections to influenza.

**Figure 7:**
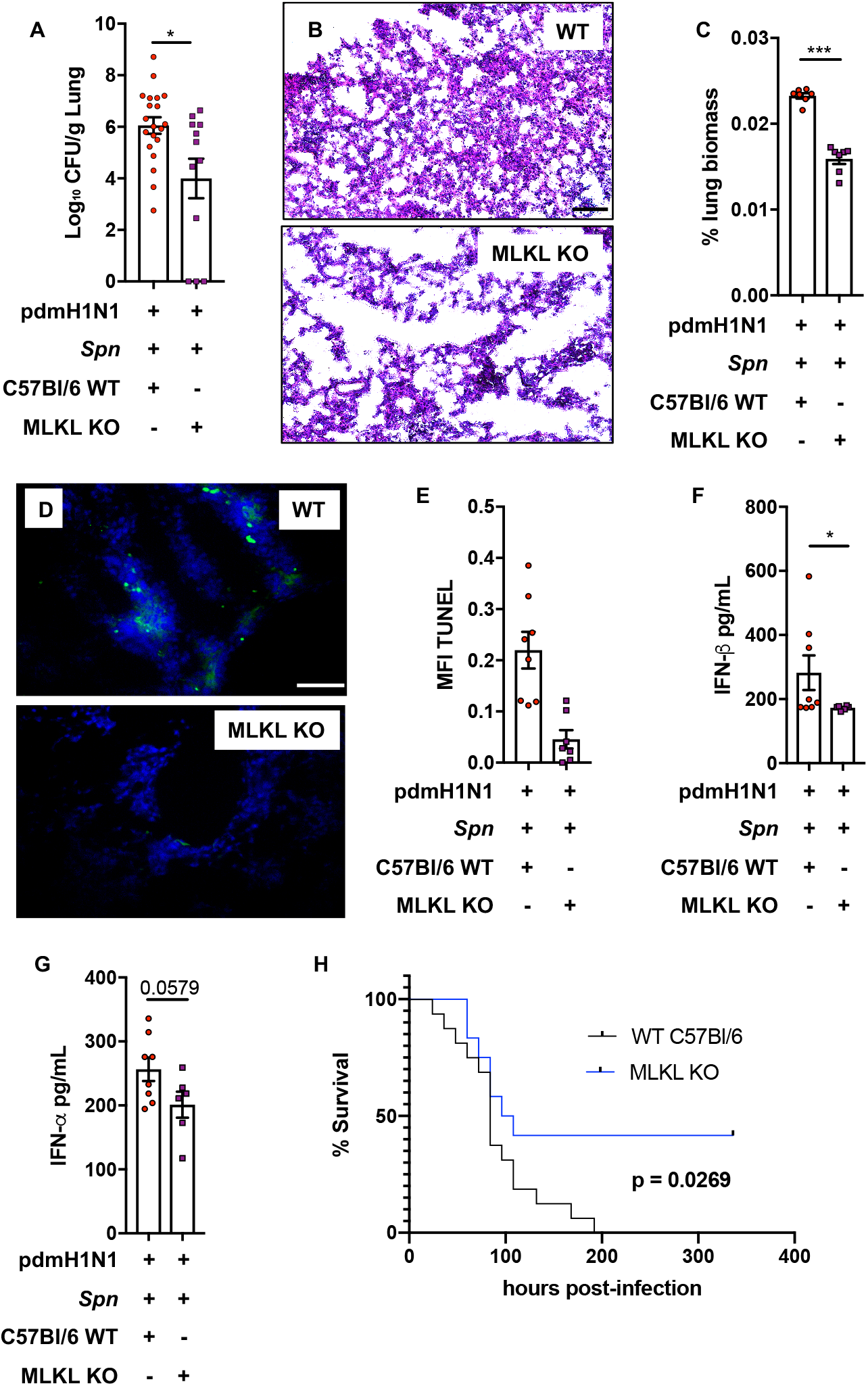
Inhibition of necroptosis reduces disease severity and tissue injury during secondary bacterial pneumonia. 8-week-old C57Bl/6 mice were intranasally infected with A/California/7/2009 (pdmH1N1) and 10 days later challenged intratracheally with *S. pneumoniae* (*Spn*). Mice were euthanized 48 hours after secondary infection (n>12 mice). **A)** Measured bacterial titers in homogenized lungs. **B)** Representative H&E staining of corresponding tissue sections. Black bar denotes 100μm. **C)** Lung consolidation in tissue sections as measured using ImageJ (white space versus lung area, separate points are the average of 3 pictures per mouse). **D)** TUNEL stain (white bar denotes 50μm) and **E)** mean fluorescent intensity of TUNEL stain quantified in lung sections. **F)** IFN-β and **G)** IFN-α levels (pg/mL) in lung homogenates. **H)** Survival of 8-week-old WT and MLKL KO-C57Bl/6 in the secondary *Spn* infection model.

## DISCUSSION

The molecular mechanisms of IAV subversion of cellular defenses and cell fate continues to be investigated (45). Only recently has it become apparent that necroptosis is essential for control of virus replication during infection (18). Herein we demonstrate that oxidative stress triggered by IAV infection plays a role in the potentiation of PFT-induced necroptosis in respiratory cells. Oxidative stress is pleiotropic and capable of oxidizing proteins and lipid membranes, damaging nucleic acid, and potentially altering cellular energy levels or ion homeostasis of the cell. The latter were shown to be triggers for non-canonical activation of necroptosis within bacteria-infected cells (16, 17, 22). During viral infection necroptosis is canonically activated through death receptor signaling and/or recognition of viral RNA and DNA by DAI (37). Whether and how the latter pathways are sensitized as result of IAV induced ROS or if an independent mechanism is responsible remains unclear, and detailed studies are now warranted to discern key similarities and differences between these events. Importantly, increased susceptibility to PFT-mediated necroptosis was still observed even when IAV replication was blocked with Pimodivir. Moreover, Tempol-mediated protection against priming for PFT killing was applicable to both the co-infection and secondary infection scenario, the latter when virus is no longer present. This suggests the mechanism responsible for IAV-mediated necroptosis potentiation is directly affected by acute intracellular ROS levels and independent of viral effectors. Our results showing no differences in caspase activation suggests the responsible mechanism is also independent of canonical apoptotic and pyroptotic pathways, although it is likely these mechanisms are contributory to overall disease and occurring in parallel during natural infection.

Sensitization to necroptosis most likely contributes to a variety of clinical problems during co- and secondary pneumonia such as acute respiratory distress syndrome and sepsis; a consequence of the enhanced level of cell death and release of pro-inflammatory alarmins. It is noteworthy that IAV has been specifically demonstrated to drive *Spn* development of otitis media (46). Critically it is unknown if other viruses enhance permissiveness for PFT-mediated necroptosis and this is an important avenue of future investigation. In support of this notion, a wide variety of viruses have been shown to induce oxidative stress in host cells by a variety of means (47). For example, respiratory syncytial virus does so by modulating levels of antioxidant enzymes (48). Thus, it is likely that this phenomena is not restricted to IAV. Our prior published work (17, 22), and that with *S. aureus* α-toxin herein, which showed a wide variety of PFT-producing bacteria can instigate necroptosis of LEC suggests viral-enhanced PFT-mediated necroptosis is not restricted to the pathogen *Spn*. What is more, this synergy may be an important contributor to enhanced disease severity at other anatomical sites where virus and bacteria can co-infect.

Finally, our results suggest that inhibition of necroptosis may be a viable therapeutic treatment during IAV mediated co- or secondary infections, although the possibility remains that necroptosis inhibition may promote viral replication during co-infection, an aspect which needs to be studied carefully (unpublished results with primary NHBEs suggest it does not). Altogether, our results provide a new molecular explanation for how influenza infection enhances permissiveness for secondary bacterial infection. We demonstrate that PFT-mediated necroptosis is enhanced as result of oxidative stress cause by prior or ongoing viral replications. Increased sensitivity to PFT-mediated necroptosis in turn worsens pulmonary damage and creates an environment that is further permissive for bacterial replication. The fact that oxidative stress induced by virus and PFT production are common across a wide range of viral and bacterial pathogens, respectively, suggests this is an important aspect of human infectious disease pathogenesis.

## Supporting information

Supplement materia

## REFERENCES

1. Morens DM, Taubenberger JK, Fauci AS. Predominant Role of Bacterial Pneumonia as a Cause of Death in Pandemic Influenza: Implications for Pandemic Influenza Preparedness. Journal of Infectious Diseases 2008; 198: 962–970.

2. Tokars JI, Olsen SJ, Reed C. Seasonal Incidence of Symptomatic Influenza in the United States. Clin Infect Dis 2018; 66: 1511–1518.

3. Clayville LR. Influenza update: a review of currently available vaccines. P T 2011; 36: 659–684.

4. van der Sluijs KF, van der Poll T, Lutter R, Juffermans NP, Schultz MJ. Bench-to-bedside review: Bacterial pneumonia with influenza - pathogenesis and clinical implications. Critical Care 2010; 14: 1–8.

5. Gill JR, Sheng ZM, Ely SF, Guinee DG, Beasley MB, Suh J, Deshpande C, Mollura DJ, Morens DM, Bray M, Travis WD, Taubenberger JK. Pulmonary pathologic findings of fatal 2009 pandemic influenza A/H1N1 viral infections. Arch Pathol Lab Med 2010; 134: 235–243.

6. Louie JK, Acosta M, Winter K, Jean C, Gavali S, Schechter R, Vugia D, Harriman K, Matyas B, Glaser CA, Samuel MC, Rosenberg J, Talarico J, Hatch D, California Pandemic Working G. Factors associated with death or hospitalization due to pandemic 2009 influenza A(H1N1) infection in California. JAMA 2009; 302: 1896–1902.

7. McCullers JA, Bartmess KC. Role of Neuraminidase in Lethal Synergism between Influenza Virus and Streptococcus pneumoniae. Journal of Infectious Diseases 2003; 187: 1000–1009.

8. Hentrich K, Löfling J, Pathak A, Nizet V, Varki A, Henriques-Normark B. Streptococcus pneumoniae Senses a Human-like Sialic Acid Profile via the Response Regulator CiaR. Cell Host & Microbe 2016; 20: 307–317.

9. Brand JD, Lazrak A, Trombley JE, Shei RJ, Adewale AT, Tipper JL, Yu Z, Ashtekar AR, Rowe SM, Matalon S, Harrod KS. Influenza-mediated reduction of lung epithelial ion channel activity leads to dysregulated pulmonary fluid homeostasis. JCI Insight 2018; 3.

10. Pettigrew MM, Marks LR, Kong Y, Gent JF, Roche-Hakansson H, Hakansson AP. Dynamic changes in the Streptococcus pneumoniae transcriptome during transition from biofilm formation to invasive disease upon influenza A virus infection. Infection and immunity 2014; 82: 4607–4619.

11. Sun K, Metzger DW. Inhibition of pulmonary antibacterial defense by interferon-[gamma] during recovery from influenza infection. Nat Med 2008; 14: 558–564.

12. Shahangian A, Chow EK, Tian X, Kang JR, Ghaffari A, Liu SY, Belperio JA, Cheng G, Deng JC. Type I IFNs mediate development of postinfluenza bacterial pneumonia in mice. The Journal of Clinical Investigation; 119: 1910–1920.

13. van der Sluijs KF, Nijhuis M, Levels JH, Florquin S, Mellor AL, Jansen HM, van der Poll T, Lutter R. Influenza-induced expression of indoleamine 2,3-dioxygenase enhances interleukin-10 production and bacterial outgrowth during secondary pneumococcal pneumonia. The Journal of infectious diseases 2006; 193: 214–222.

14. Vandenabeele P, Galluzzi L, Vanden Berghe T, Kroemer G. Molecular mechanisms of necroptosis: an ordered cellular explosion. Nat Rev Mol Cell Biol 2010; 11: 700–714.

15. Moreno-Gonzalez G, Vandenabeele P, Krysko DV. Necroptosis: A Novel Cell Death Modality and Its Potential Relevance for Critical Care Medicine. Am J Respir Crit Care Med 2016; 194: 415–428.

16. Gonzalez-Juarbe N, Bradley KM, Riegler AN, Reyes LF, Brissac T, Park SS, Restrepo MI, Orihuela CJ. Bacterial Pore-Forming Toxins Promote the Activation of Caspases in Parallel to Necroptosis to Enhance Alarmin Release and Inflammation During Pneumonia. Sci Rep 2018; 8: 5846.

17. Gonzalez-Juarbe N, Bradley KM, Shenoy AT, Gilley RP, Reyes LF, Hinojosa CA, Restrepo MI, Dube PH, Bergman MA, Orihuela CJ. Pore-forming toxin-mediated ion dysregulation leads to death receptor-independent necroptosis of lung epithelial cells during bacterial pneumonia. Cell Death Differ 2017; 24: 917–928.

18. Nogusa S, Thapa RJ, Dillon CP, Liedmann S, Oguin TH, 3rd, Ingram JP, Rodriguez DA, Kosoff R, Sharma S, Sturm O, Verbist K, Gough PJ, Bertin J, Hartmann BM, Sealfon SC, Kaiser WJ, Mocarski ES, Lopez CB, Thomas PG, Oberst A, Green DR, Balachandran S. RIPK3 Activates Parallel Pathways of MLKL-Driven Necroptosis and FADD-Mediated Apoptosis to Protect against Influenza A Virus. Cell Host Microbe 2016; 20: 13–24.

19. Wang Y, Hao Q, Florence JM, Jung BG, Kurdowska AK, Samten B, Idell S, Tang H. Influenza Virus Infection Induces ZBP1 Expression and Necroptosis in Mouse Lungs. Front Cell Infect Microbiol 2019; 9: 286.

20. Thapa RJ, Ingram JP, Ragan KB, Nogusa S, Boyd DF, Benitez AA, Sridharan H, Kosoff R, Shubina M, Landsteiner VJ, Andrake M, Vogel P, Sigal LJ, tenOever BR, Thomas PG, Upton JW, Balachandran S. DAI Senses Influenza A Virus Genomic RNA and Activates RIPK3-Dependent Cell Death. Cell Host Microbe 2016; 20: 674–681.

21. Gonzalez-Juarbe N, Bradley KM, Shenoy AT, Gilley RP, Reyes LF, Hinojosa CA, Restrepo MI, Dube PH, Bergman MA, Orihuela CJ. Pore-forming toxin-mediated ion dysregulation leads to death receptor-independent necroptosis of lung epithelial cells during bacterial pneumonia. Cell Death Differ 2017.

22. Gonzalez-Juarbe N, Gilley RP, Hinojosa CA, Bradley KM, Kamei A, Gao G, Dube PH, Bergman MA, Orihuela CJ. Pore-Forming Toxins Induce Macrophage Necroptosis during Acute Bacterial Pneumonia. PLoS Pathog 2015; 11: e1005337.

23. Fulcher ML, Gabriel S, Burns KA, Yankaskas JR, Randell SH. Well-differentiated human airway epithelial cell cultures. Methods Mol Med 2005; 107: 183–206.

24. Tettelin H, Nelson KE, Paulsen IT, Eisen JA, Read TD, Peterson S, Heidelberg J, DeBoy RT, Haft DH, Dodson RJ, Durkin AS, Gwinn M, Kolonay JF, Nelson WC, Peterson JD, Umayam LA, White O, Salzberg SL, Lewis MR, Radune D, Holtzapple E, Khouri H, Wolf AM, Utterback TR, Hansen CL, McDonald LA, Feldblyum TV, Angiuoli S, Dickinson T, Hickey EK, Holt IE, Loftus BJ, Yang F, Smith HO, Venter JC, Dougherty BA, Morrison DA, Hollingshead SK, Fraser CM. Complete genome sequence of a virulent isolate of Streptococcus pneumoniae. Science 2001; 293: 498–506.

25. Lizcano A, Chin T, Sauer K, Tuomanen EI, Orihuela CJ. Early biofilm formation on microtiter plates is not correlated with the invasive disease potential of Streptococcus pneumoniae. Microbial pathogenesis 2010; 48: 124–130.

26. Zafar MA, Wang Y, Hamaguchi S, Weiser JN. Host-to-Host Transmission of Streptococcus pneumoniae Is Driven by Its Inflammatory Toxin, Pneumolysin. Cell Host Microbe 2017; 21: 73–83.

27. Brown AO, Mann B, Gao G, Hankins JS, Humann J, Giardina J, Faverio P, Restrepo MI, Halade GV, Mortensen EM, Lindsey ML, Hanes M, Happel KI, Nelson S, Bagby GJ, Lorent JA, Cardinal P, Granados R, Esteban A, LeSaux CJ, Tuomanen EI, Orihuela CJ. Streptococcus pneumoniae translocates into the myocardium and forms unique microlesions that disrupt cardiac function. PLoS Pathog 2014; 10: e1004383.

28. Murphy JM, Czabotar PE, Hildebrand JM, Lucet IS, Zhang JG, Alvarez-Diaz S, Lewis R, Lalaoui N, Metcalf D, Webb AI, Young SN, Varghese LN, Tannahill GM, Hatchell EC, Majewski IJ, Okamoto T, Dobson RC, Hilton DJ, Babon JJ, Nicola NA, Strasser A, Silke J, Alexander WS. The pseudokinase MLKL mediates necroptosis via a molecular switch mechanism. Immunity 2013; 39: 443–453.

29. Gonzalez-Juarbe N, Mares CA, Hinojosa CA, Medina JL, Cantwell A, Dube PH, Orihuela CJ, Bergman MA. Requirement for Serratia marcescens cytolysin in a murine model of hemorrhagic pneumonia. Infect Immun 2015; 83: 614–624.

30. Saxena RK, Gilmour MI, Hays MD. Isolation and quantitative estimation of diesel exhaust and carbon black particles ingested by lung epithelial cells and alveolar macrophages in vitro. Biotechniques 2008; 44: 799–805.

31. Gilley RP, Gonzalez-Juarbe N, Shenoy AT, Reyes LF, Dube PH, Restrepo MI, Orihuela CJ. Infiltrated Macrophages Die of Pneumolysin-Mediated Necroptosis following Pneumococcal Myocardial Invasion. Infect Immun 2016; 84: 1457–1469.

32. Schindelin J, Rueden CT, Hiner MC, Eliceiri KW. The ImageJ ecosystem: An open platform for biomedical image analysis. Mol Reprod Dev 2015; 82: 518–529.

33. Riegler AN, Brissac T, Gonzalez-Juarbe N, Orihuela CJ. Necroptotic Cell Death Promotes Adaptive Immunity Against Colonizing Pneumococci. Front Immunol 2019; 10: 615.

34. McCullers JA, Hayden FG. Fatal influenza B infections: time to reexamine influenza research priorities. J Infect Dis 2012; 205: 870–872.

35. McCullers JA, Tuomanen EI. Molecular pathogenesis of pneumococcal pneumonia. Front Biosci 2001; 6: D877–D889.

36. Hoffmann J, Machado D, Terrier O, Pouzol S, Messaoudi M, Basualdo W, Espinola EE, Guillen RM, Rosa-Calatrava M, Picot V, Benet T, Endtz H, Russomando G, Paranhos-Baccala G. Viral and bacterial co-infection in severe pneumonia triggers innate immune responses and specifically enhances IP-10: a translational study. Sci Rep 2016; 6: 38532.

37. Upton JW, Shubina M, Balachandran S. RIPK3-driven cell death during virus infections. Immunol Rev 2017; 277: 90–101.

38. Morris DE, Cleary DW, Clarke SC. Secondary Bacterial Infections Associated with Influenza Pandemics. Frontiers in microbiology 2017; 8: 1041–1041.

39. Liu M, Chen F, Liu T, Chen F, Liu S, Yang J. The role of oxidative stress in influenza virus infection. Microbes Infect 2017; 19: 580–586.

40. Brissac T, Shenoy AT, Patterson LA, Orihuela CJ. Cell invasion and pyruvate oxidase derived H2O2 are critical for Streptococcus pneumoniae mediated cardiomyocyte killing. Infect Immun 2017.

41. Takahashi K, Furuta Y, Fukuda Y, Kuno M, Kamiyama T, Kozaki K, Nomura N, Egawa H, Minami S, Shiraki K. In vitro and in vivo activities of T-705 and oseltamivir against influenza virus. Antivir Chem Chemother 2003; 14: 235–241.

42. Byrn RA, Jones SM, Bennett HB, Bral C, Clark MP, Jacobs MD, Kwong AD, Ledeboer MW, Leeman JR, McNeil CF, Murcko MA, Nezami A, Perola E, Rijnbrand R, Saxena K, Tsai AW, Zhou Y, Charifson PS. Preclinical activity of VX-787, a first-in-class, orally bioavailable inhibitor of the influenza virus polymerase PB2 subunit. Antimicrob Agents Chemother 2015; 59: 1569–1582.

43. Helbock HJ, Beckman KB, Ames BN. 8-Hydroxydeoxyguanosine and 8-hydroxyguanine as biomarkers of oxidative DNA damage. Methods Enzymol 1999; 300: 156–166.

44. Kruman I, Bruce-Keller AJ, Bredesen D, Waeg G, Mattson MP. Evidence that 4-hydroxynonenal mediates oxidative stress-induced neuronal apoptosis. J Neurosci 1997; 17: 5089–5100.

45. Yeganeh B, Rezaei Moghadam A, Tran AT, Rahim MN, Ande SR, Hashemi M, Coombs KM, Ghavami S. Asthma and influenza virus infection:focusing on cell death and stress pathways in influenza virus replication. Iran J Allergy Asthma Immunol 2013; 12: 1–17.

46. Tong HH, Fisher LM, Kosunick GM, DeMaria TF. Effect of adenovirus type 1 and influenza A virus on Streptococcus pneumoniae nasopharyngeal colonization and otitis media in the chinchilla. Ann Otol Rhinol Laryngol 2000; 109: 1021–1027.

47. Schwarz KB. Oxidative stress during viral infection: a review. Free Radic Biol Med 1996; 21: 641–649.

48. Hosakote YM, Liu T, Castro SM, Garofalo RP, Casola A. Respiratory syncytial virus induces oxidative stress by modulating antioxidant enzymes. Am J Respir Cell Mol Biol 2009; 41: 348–357.

