## Supplement materia for "Influenza-induced oxidative stress sensitizes lung cells to bacterial toxin-mediated necroptosis"

**Online Data Supplement:**

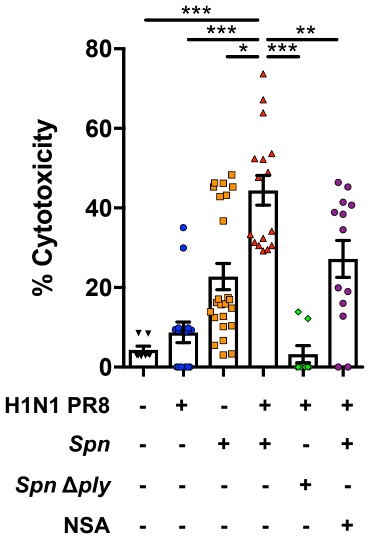


**Figure E1: IAV infection augments pneumolysin-mediated necroptosis.**

Cytotoxicity was assessed using the LDH assay. A549 cells were infected with PR8 H1N1 at a MOI 2 for 2 hours and subsequently challenged with wildtype *S. pneumoniae* (*Spn*) or Ply deficient mutant (*Spn* Δ*ply*) at a MOI of 10 for 4 hours. When noted, cells were pretreated with necrosulfonamide (NSA, 10μM) for 1 hour prior to bacterial challenge.

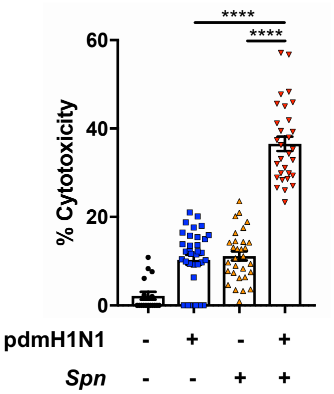


**Figure E2: IAV infection augments *Spn*-mediated cytotoxicity in a macrophage cell line.** Cytotoxicity was assessed using the LDH assay. MH-S murine alveolar macrophages were infected with pdmH1N1 at a MOI 2 for 2 hours. Cells were subsequently challenged with *S. pneumoniae* (*Spn*) at a MOI of 10 for 4 hours.

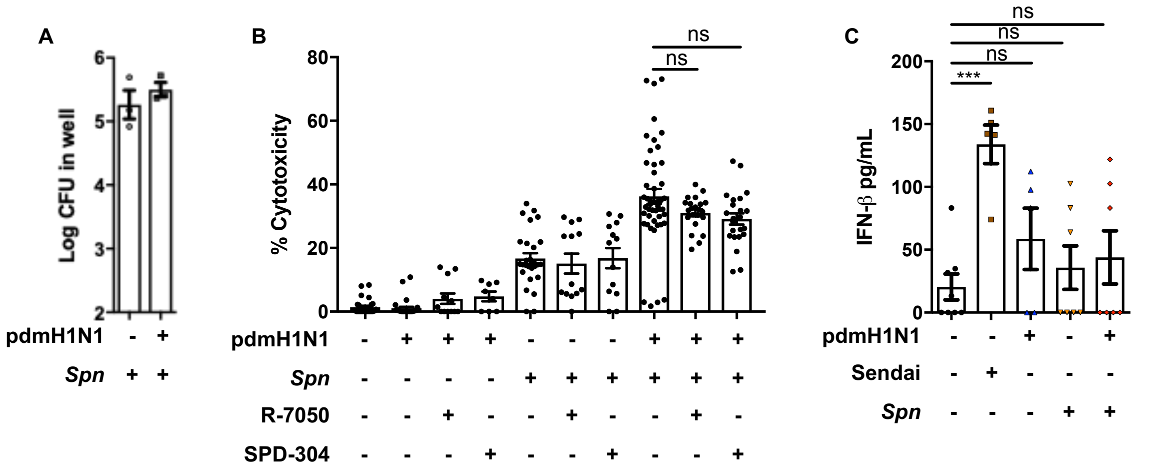


**Figure E3: Primary IAV infection does not affect bacterial titers and potentiation of necroptosis does not require inflammatory signals *in vitro*. A)** Shown is the amount of *Spn* recovered from 24-well A549 monolayers following pdmH1N1/*Spn* co-infection. **B)** Cytotoxicity was assessed using the LDH assay. A549 cells were infected with pdmH1N1 at a MOI 2 for 2 hours. Cells were then challenged with *Spn* at an MOI of 10 for 4 hours. When indicated, cells were also pretreated with R-7050, a TNFr1 inhibitor (R-7050, 10μM) or SPD-304 (10μM), a TNF-α inhibitor. **C)** Levels of IFN-β detected in supernatants from A549 cells challenged with Sendai virus (Sendai), pdmH1N1, *Spn* for 4 hours, or pdmH1N1 followed with *Spn* for 4 hours was measured by ELISA.

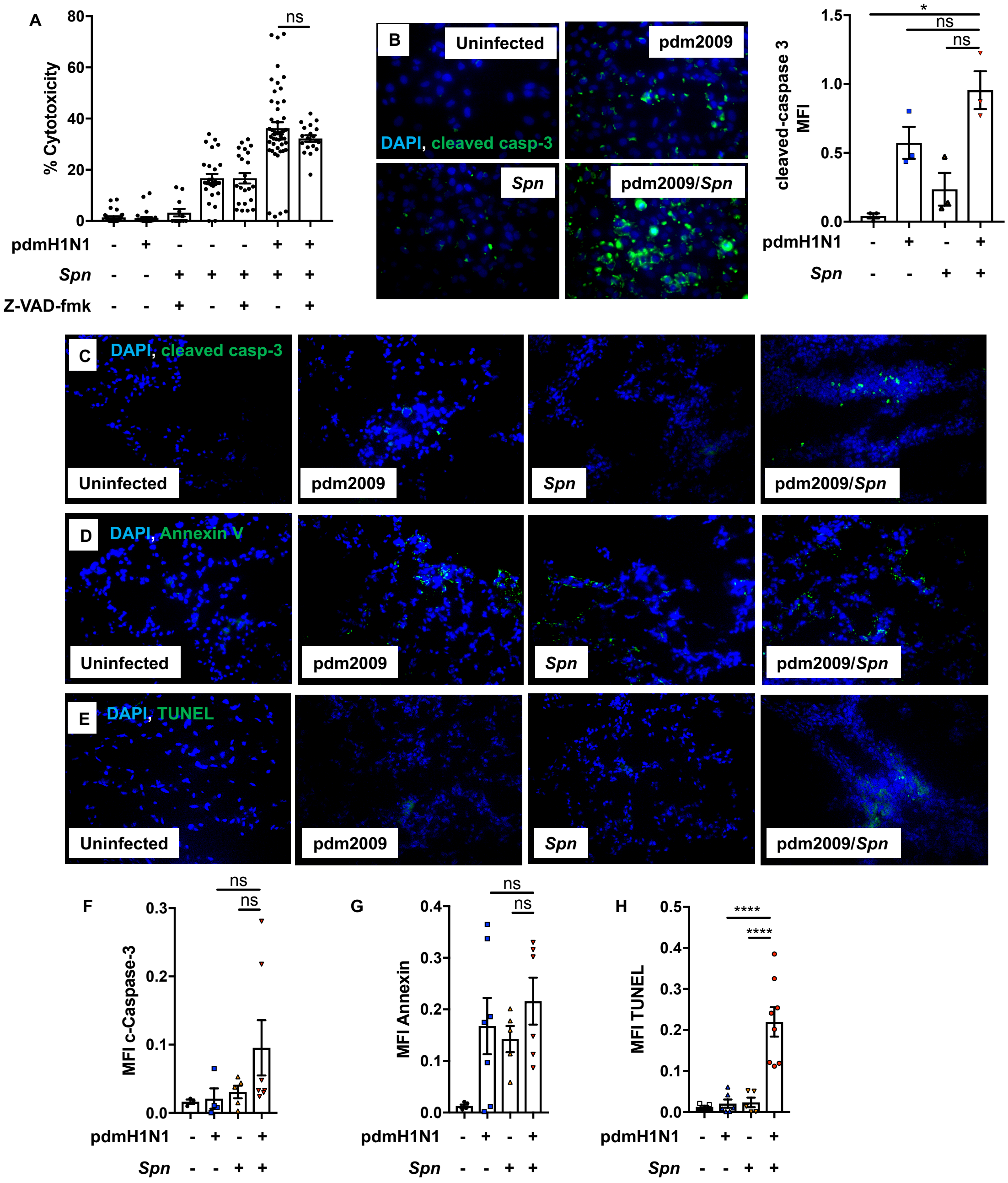


**Figure E4: Primary IAV infection does not potentiate apoptosis. A)** Cytotoxicity was assessed using the LDH assay. A549 cells were infected with pdmH1N1 at a MOI 2 for 2 hours, subsequently, cells were challenged with *S. pneumoniae* strain TIGR4 (*Spn*) at an MOI of 10 for 4 hours. When noted, cells were pretreated with general caspase inhibitor Z-VAD-fmk (100μM) for 1 hour prior to bacterial challenge. **B)** Shown are representative images and corresponding mean fluorescent intensity (MFI) of immunofluorescent stained A549 cells for cleaved-caspase 3 (green) in our co-infection model. **C-E)** Shown are representative images of whole lung sections from 8-week-old C57Bl/6 mice intranasally infected with pdmH1N1 for 10 days and subsequently challenged intratracheally with *Spn* for 48 prior to sacrifice. Shown sections were stained for **C)** cleaved caspase-3, **D)** Annexin-V, and **E)** TUNEL, and **F-H**) corresponding MFI measurements were calculated.

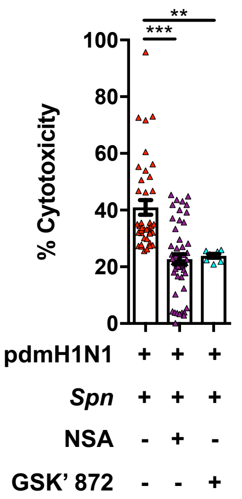


**Figure E5: IAV infection augments pneumolysin mediated RIPK3-MLKL dependent necroptosis.** Cytotoxicity was assessed using the LDH assay. A549 cells were infected with pdmH1N1 at a MOI 2 for 2 hours, subsequently, cells were challenged with *S. pneumoniae* strain TIGR4 (*Spn*) at a MOI of 10 for 4 hours. When noted, cells were pretreated with necrosulfonamide (NSA, 10µM) or GSK' 872 (GSK, 10µM) for 1 hour prior to bacterial challenge.

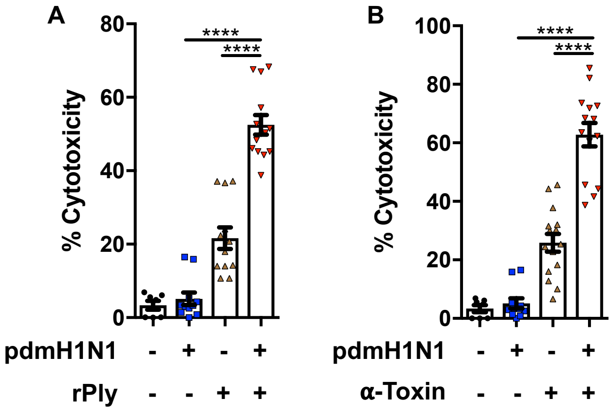


**Figure E6: IAV leads to sensitization of pore-forming toxin mediated necroptosis.** **A)** Cytotoxicity was assessed using the LDH assay. A549 cells were infected with pdmH1N1 at an MOI 2 for 2 hours then challenged with **A)** recombinant pneumolysin (rPly, 0.1 µg/ml) or **B)** recombinant alpha-toxin (α-Toxin, 0.6 µg/ml) for two more hours.

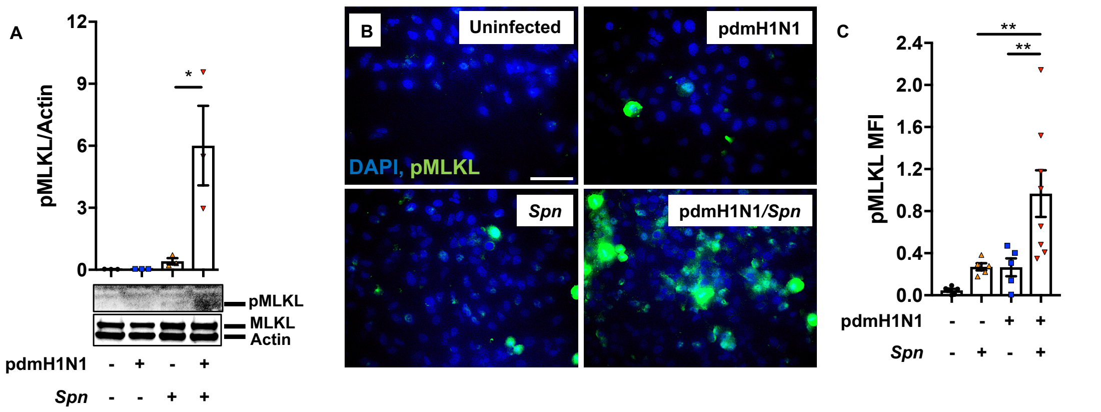


**Figure E7: MLKL is active during IAV-mediated potentiation of necroptosis. A)** Shown immunoblots for p-MLKL, MLKL and actin of A549 cells following our pdmH1N1/*Spn* co-infection model. **B)** Immunofluorescent staining for p-MLKL and **C)** mean fluorescent intensity of p-MLKL in A549 cells following their co-infection.

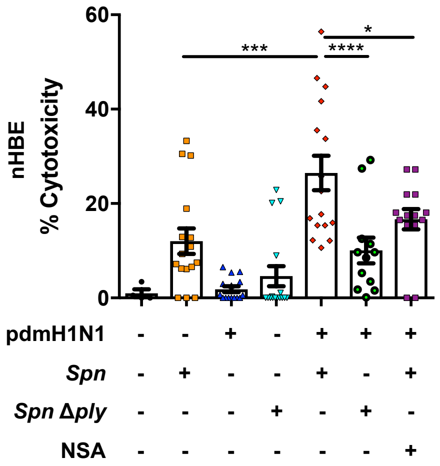


**Figure E8: IAV infection promotes pneumolysin mediated necroptosis in primary human normal bronchial epithelial cells.** Cytotoxicity was assessed using the LDH assay. *Ex vivo* cultured primary lung epithelial cells were infected with pdmH1N1 at a MOI of 2 for 2 hours. Cells were then challenged with *Spn* or Ply deficient mutant (*Spn* Δply) at a MOI of 10 for 4 additional hours. When noted, cells were treated with necrosulfonamide (NSA, 10μM) prior to bacterial challenge.

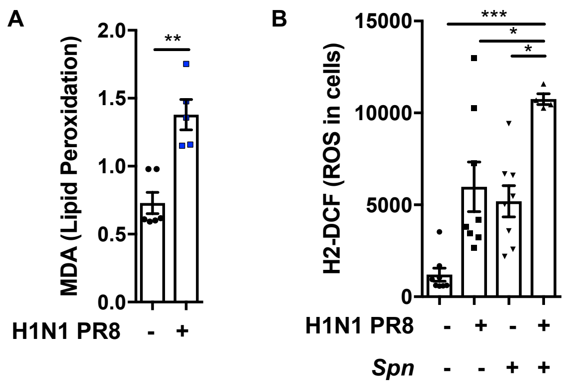


**Figure E9: IAV-mediated oxidative stress is required for the potentiation of pneumolysin mediated necroptosis. A)** Lipid peroxidation levels 4-hours after challenge with PR8 H1N1 was quantitated by MDA. **B)** Levels of cellular ROS in A549 cells infected with PR8 H1N1 at a MOI 2 for 2 hours then challenged with *Spn* at a MOI of 10 for 2 hours was assessed by H2-DCF staining.

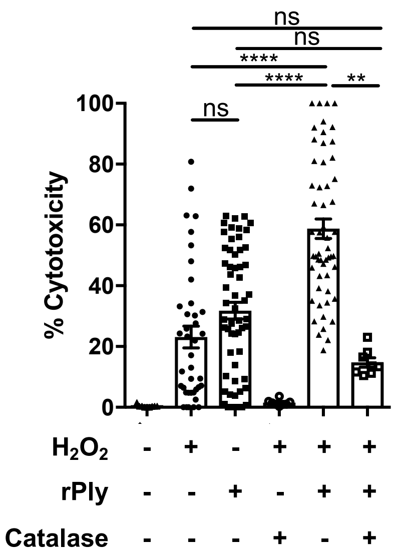


**Figure E10: Hydrogen peroxide promotes pneumolysin mediated necroptosis.** Cytotoxicity was assessed using the LDH assay. Cells were treated with combinations of hydrogen peroxide (100 µM), catalase (100 µM) for 2 hours and then challenged with rPly (0.1 µg/ml).

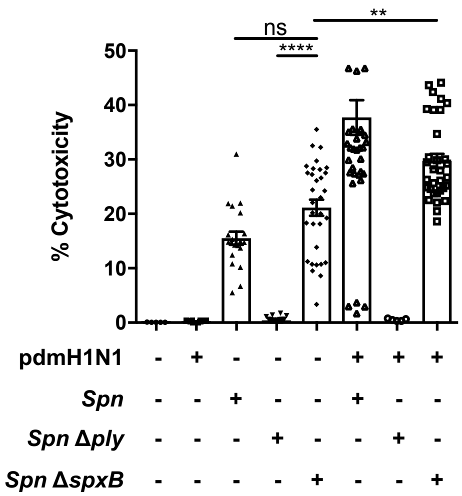


**Figure E11: Bacterial produced hydrogen peroxide is not required for potentiation of toxin mediated necroptosis during co-infection.** Cytotoxicity was assessed using the LDH assay. A549 cells were infected with pdmH1N1 at MOI of 2 for 2 hours then challenged with *Spn*, *Spn* Δ*ply*, or *Spn* Δ*spxB* at MOI of 10 for 4 hours.

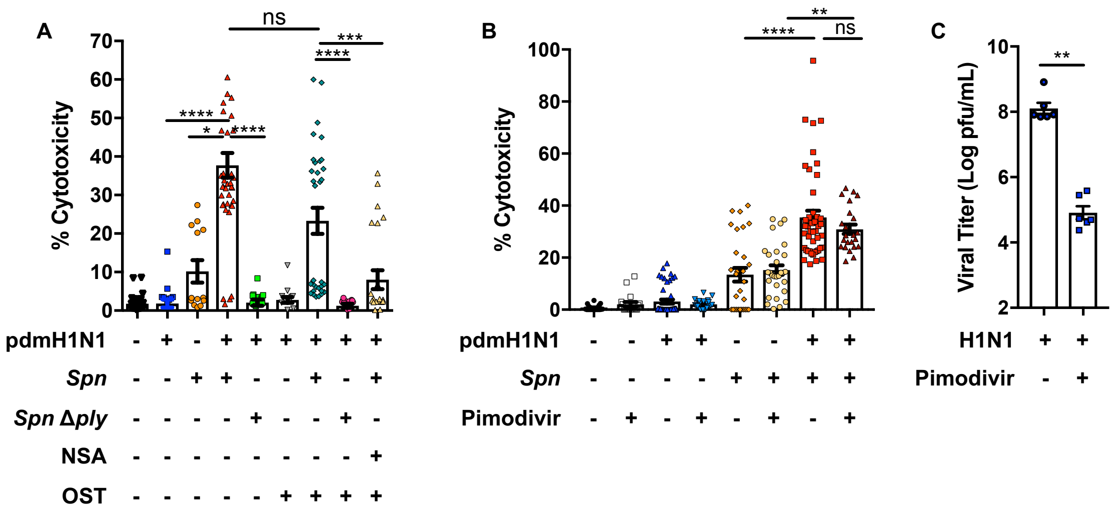


**Figure E12: Viral release and replication does not signal for influenza augmentation of PFT-mediated necroptosis.** Cytotoxicity was assessed using the LDH assay. A549 cells were infected with pdmH1N1at a MOI 2 for 2 hours and subsequently challenged with *Spn* or *Spn* Δ*ply*. **A)** When noted cells were treated with oseltamivir (one hour-post pdmH1N1 infection) or treated with NSA prior to bacterial challenge. **B)** In similar experiments cells were treated with Pimodivir (0.5µM, two-hours post initial infection) for 12 hours and subsequently challenged with *Spn*. **C)** Viral titers measured after treatment of cells with Pimodivir (0.5µM) 12 hours prior and 2-hour post pdmH1N1 initial infection.

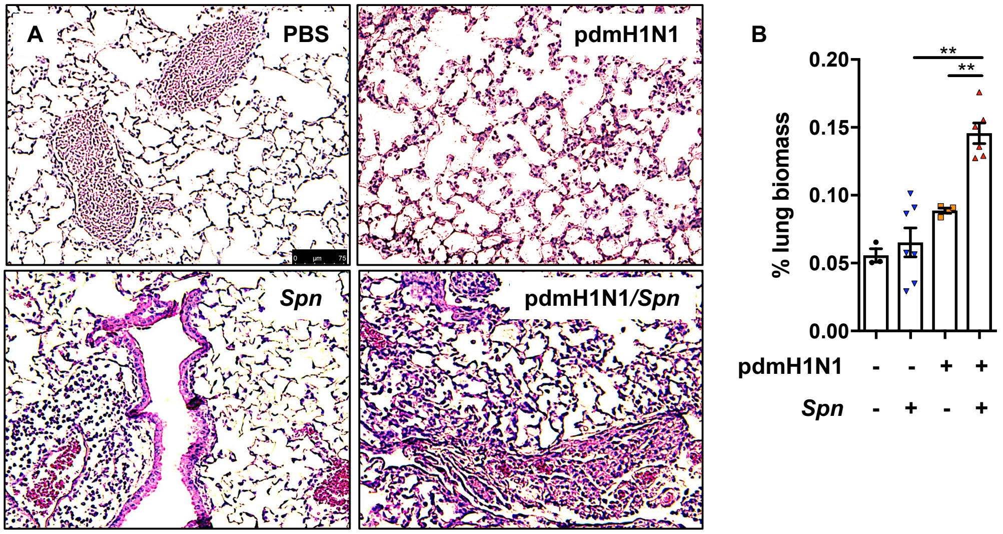


**Figure E13: Influenza promotes tissue inflammation and consolidation during secondary bacterial challenge.** 8-week-old C57Bl/6 mice were intranasally infected with pdmH1N1 for 10 days and subsequently challenged intratracheally with *Spn.* Mice were euthanized 48 hours' post-secondary infection (n=3-7 mice). **A)** Show are hematoxylin and eosin stained representative lung tissue sections. Black bar denotes 75µm. **B)** Lung consolidation was determined by measuring the stained biomass within the same corresponding lung space using ImageJ. Each point corresponds to the average calculated using 3 pictures per mouse.

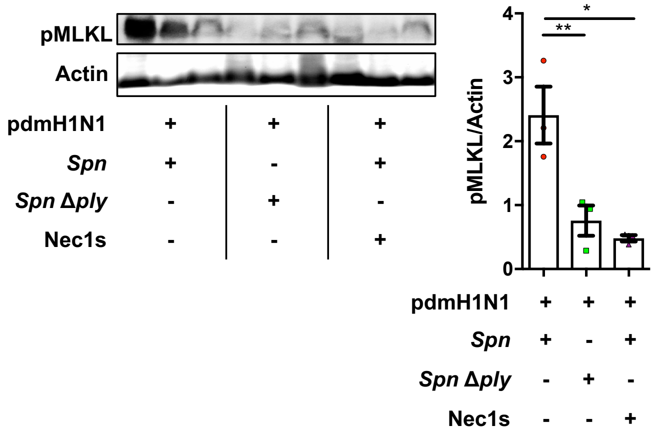


**Figure E14: Influenza infection promotes pneumolysin mediated necroptosis.** Shown is an Immunoblot for p-MLKL and actin of pdmH1N1 infected mice, challenged with *Spn*, *Spn* Δ*ply* or *Spn* with subsequent Nec1s treatment and its densitometry quantification.

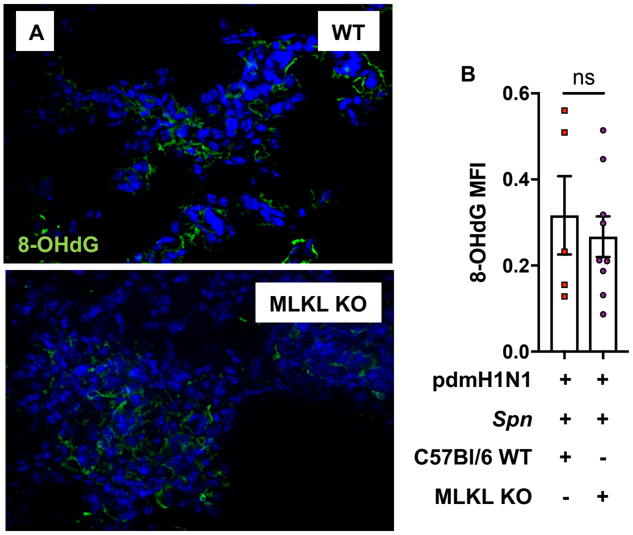


**Figure E15: Necroptosis deficiency does not affect influenza mediated oxidative stress mediated damage.** 8-week-old WT and MLKL KO C57Bl/6 mice were intranasally infected with pdmH1N1 for 10 days, subsequently challenged intratracheally with *Spn,* and euthanized 48 hours post-bacterial challenge (n=5-9 mice). (**A-B**) Immunofluorescent staining and mean fluorescent intensity for oxidative stress DNA damage marker 8-Hydroxydeoxyguanosine (8-OHdG).
